# Structural Plasticity of the Membrane-Bound Protein Degradation Assembly Supports Bacterial Adaptation to Stress

**DOI:** 10.1101/2025.07.21.662073

**Authors:** Naseer Iqbal, Sandro Keller, Alireza Ghanbarpour

## Abstract

Protein degradation by AAA+ proteases is essential for bacterial adaptation to environmental stress. The membrane-bound AAA+ protease FtsH forms a large inner-membrane complex with the SPFH (Stomatin, Prohibitin, Flotillin, HflK/C) family transmembrane proteins HflK and HflC, playing a key role in bacterial recovery from aminoglycoside antibiotic stress. Recent structural studies have revealed both open, asymmetric and closed, symmetric conformations of the HflK/C assembly under different sample-preparation conditions, suggesting two distinct models for how this complex modulates FtsH proteolysis. To determine which conformation reflects the biologically active state, we engineered a disulfide-crosslinked HflK/C variant to stabilize the closed conformation and resolved its structure using high-resolution cryo-EM. Phenotypic assays showed that cells expressing either this stabilized, closed HflK/C variant or an HflK/C mutant that disrupts interactions with FtsH exhibit significantly impaired growth under aminoglycoside stress. Surprisingly, the cryo-EM structure of the FtsH•HflK/C complex from cells challenged with the aminoglycoside antibiotic tobramycin revealed a novel HflK/C arrangement, characterized by two openings on opposite sides that may facilitate substrate access to FtsH during proteotoxic stress. Together, our results suggest that both the dynamic open conformation of HflK/C and its specific interactions with FtsH are critical for adaptation to aminoglycoside-induced stress. Given the conserved structural and functional features of SPFH family members, our findings may offer a broader framework for understanding how this protein family operates under both basal and stress conditions.

## INTRODUCTION

Dynamic protein degradation allows organisms, from bacteria to humans, to regulate their proteome in response to diverse cellular signals^1^. In bacteria, this process is primarily mediated by five major AAA (ATPases Associated with diverse Cellular Activities) proteases, with FtsH being the only membrane-spanning member, which degrades both soluble and membrane-associated proteins^2-3^. Structurally, an FtsH monomer consists of a transmembrane helix, a periplasmic domain, a second transmembrane helix, and a cytoplasmic AAA+ module, followed by a zinc-dependent peptidase domain^4-5^.

The AAA+ domain recognizes substrates via a degron sequence and uses ATP hydrolysis to generate the mechanical force needed for substrate unfolding and translocation into the peptidase chamber for degradation^6-7^. In *Escherichia coli*, FtsH is essential as it plays a pivotal role in lipid metabolism by degrading key enzymes involved in lipopolysaccharide (LPS) biosynthesis, including LpxC and KdtA^8-9^. This degradation helps balance phospholipid and LPS synthesis, regulating outer membrane permeability and ensuring proper cell-envelope biogenesis^8^. Additionally, FtsH degrades unpartnered SecY and additional misfolded or damaged membrane proteins, helping maintain integrity of protein translocation across the inner membrane^10^. In eukaryotes, mitochondrial FtsH homologs are essential for protein quality control, with dysregulation linked to neurodegenerative and metabolic diseases^11-13^.

FtsH associates with the single-transmembrane proteins HflK and HflC to form a massive complex in the bacterial inner membrane^14^. HflK and HflC belong to the SPFH (Stomatin, Prohibitin, Flotillin, HflK/C) protein family, which is conserved across bacteria and eukaryotes and is known to organize membrane microdomains and regulate membrane-associated processes^15-17^. Although the HflK/C complex is dispensable under normal conditions, it becomes essential during stress, such as exposure to aminoglycoside antibiotics or oxidative conditions^18^, which result in the accumulation of unfolded or damaged proteins that must be cleared in an FtsH-and HflK/C-dependent manner^19^. Deletion of *hflK* and *hflC* also impairs bacterial aerobic respiration^20^, resembling mitochondrial defects caused by prohibitin inactivation in yeast and mammalian cells^20-21^.

Previous studies involving overproduction of the complex demonstrated that HflK/C forms a closed cage around four copies of FtsH^22-23^. Based on this structure, the authors proposed a model in which HflK/C cage inhibits FtsH-mediated proteolysis of membrane-protein substrates^22-23^. Through extraction of the endogenous complex using both detergent-based and detergent-free methods, we have previously revealed a markedly different arrangement: two hexameric FtsH proteases embedded within an open, asymmetric, nautilus-like HflK/C assembly^5^. Based on structural and proteomic analyses, we proposed that the HflK/C assembly modulates FtsH activity both positively and negatively^5^.

Herein, to elucidate how different conformations of the complex contribute to bacterial recovery from stress, we engineered a disulfide-crosslinked variant of HflK/C that stably adopts a closed conformation. Compared to the wild-type HflK/C, phenotypic assays revealed that the closed-conformation variant impaired bacterial growth during treatment with tobramycin, an aminoglycoside antibiotic. Cryo-EM analysis of native complexes extracted from tobramycin-treated cells revealed a new conformation of the FtsH•HflK/C membrane complex, featuring two openings positioned adjacent to both FtsH hexamers. Together, our structural and phenotypic data support a model in which conformational flexibility of the HflK/C complex is essential for regulating FtsH activity and facilitating bacterial adaptation to antibiotic stress.

## RESULTS

### Designing disulfide bond cross-links between HflK and HflC to stabilize the closed conformation

To investigate the biological significance of the open, nautilus-like assembly and its opening in regulating membrane-protein degradation by FtsH, we engineered a disulfide bond crosslink between HflK and HflC to stabilize the closed HflK/C assembly.

Guided by our previous structural data (PDB ID: 9CZ2), we introduced cysteine substitutions at alanine residues 270 and 283 in HflK, and at residues 233 and 264 in HflC, with the goal of generating two disulfide bonds—each connecting one HflK subunit to two adjacent HflC subunits, and vice versa (Fig. 1a). These positions were selected for three reasons. First, the “hat” portion of the structure is relatively rigid, as it shows limited conformational variability across available structures^5, 22-23^. This structural rigidity enhances the likelihood of successful disulfide bond formation, as the targeted residues are not subject to significant positional shifts (Fig. 1a). Second, the opening observed in our previous structure (PDB ID: 9CZ2) originates from the coiled-coil domain of four HflK/C subunits (Fig. 1a, close-up view). The last modeled residues in this flexible region still include the engineered cysteine mutations. Therefore, introducing disulfide bonds at these sites was expected to restrict conformational flexibility and stabilize the closed conformation. Third, because these residues are located in the periplasmic space, its oxidative environment naturally promotes disulfide bond formation.

**Figure 1.**
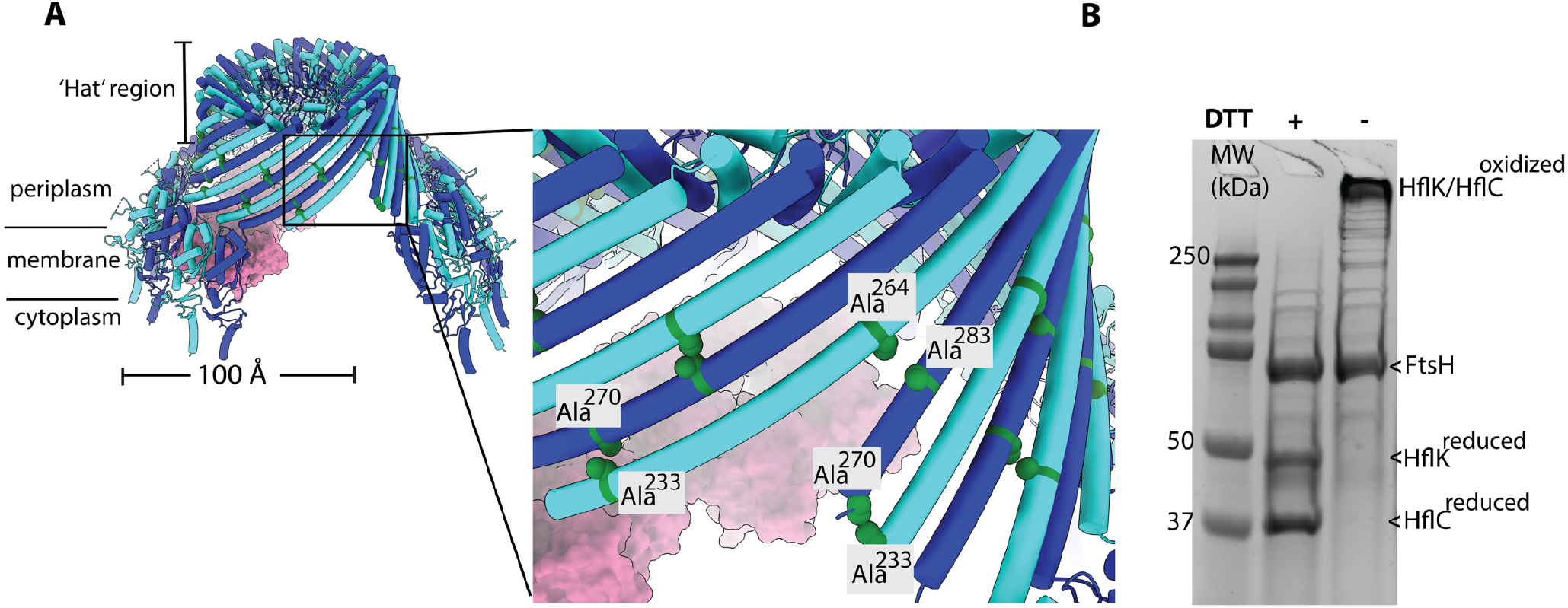
Designing disulfide bonds between HflK and HflC to stabilize the closed conformation of the HflK/C membrane assembly. (**A**) Positions of cysteine substitutions in HflK (Ala^270^ and Ala^283^) and HflC (Ala^233^ and Ala^264^). The mutations are designed in the ‘hat’ region of the structure and allow each HflK to be crosslinked to two neighboring HflC subunits, and vice versa. HflK and HflC are shown in blue and cyan, respectively, in cartoon representation. (**B**) SDS-PAGE analysis of affinity-purified FtsH•HflK/C (right lane) under reducing and non-reducing conditions, with molecular weight standards in the left lane. Under reducing conditions, distinct bands corresponding to HflK and HflC are observed; under oxidizing conditions, these merge into a high-molecular-weight band that does not migrate through the gel, indicating disulfide bond formation under non-reducing condition.

### Validation of the disulfide cross-linked HflK/C (HflK/C^SS^) and its interaction with FtsH

We expressed the disulfide cross-linked HflK/C mutant (HflK/C^**SS**^) using a low-copy-number plasmid under the control of a sodium propionate–titratable promoter in *E. coli* BL21 cells lacking the endogenous *hflK/C* genes. Additionally, these cells harbored a chromosomally encoded FLAG tag at the C-terminus of FtsH to facilitate complex pull-down^5^. Following expression of HflK/C^**SS**^, the complex was affinity-purified using M2-FLAG resin. SDS-PAGE analysis of the eluate under reducing conditions confirmed successful co-purification of FtsH along with two additional protein bands corresponding to the sizes of HflK and HflC (Fig. 1b). Notably, under nonreducing conditions, the HflK and HflC bands merged into a single high–molecular-weight species that failed to migrate into the gel, consistent with the formation of a larger HflK/C assembly stabilized by disulfide bonds. The ratio of band intensities on SDS-PAGE for the HflK and HflC mutants relative to FtsH was comparable to that of wild-type HflK/C (HflK/C^**WT**^), indicating that disulfide bond formation did not affect the interaction between FtsH and HflK/C.

### Cryo-EM structure of the closed HflK/C assembly

To verify that cross-linking between HflK and HflC did not cause aberrant aggregation of the massive HflK/C complex, we prepared cryo-EM grids of the FtsH•HflK/C^**SS**^ complex solubilized in n-dodecyl β-D-maltoside (DDM) and glyco-diosgenin (GDN) (Figs. 2 and S1–S3). Initial screening revealed 2D class averages consistent with a fully closed conformation of the HflK/C^**SS**^ cage under non-reducing conditions (Fig. 2A). We also prepared another cryo-EM grid using the complex after incubation with DTT for 1 hour at 30 °C (Fig. 2B). Screening of this grid revealed an open, nautilus-like assembly resembling wild-type HflK/C (PDB ID: 9CZ2)^5^, indicating that the alanine-to-cysteine mutations *per se* did not disrupt the open, nautilus-like HflK/C conformation and that disulfide bond formation enforced the closed state. Following 3D reconstruction of the oxidized FtsH•HflK/C^**SS**^ complex, we determined the cryo-EM structure of HflK/C^**SS**^ at a global resolution of 2.9 Å in DDM using C_1_ symmetry (Fig. 2, Figs. S1–S3, Table 1). The density map resolved 24 copies of HflK/C, with the exception of the Nand C-termini of HflK, which are predicted to be unstructured^5^, and the transmembrane regions of HflK/C and FtsH, which showed poor density (Fig. 2 and Table 1). Although the entire complex was pulled down via FtsH, FtsH itself was not seen in the final 3D reconstruction—likely due to the high symmetry of the closed HflK/C^**SS**^ structure, where each of the 12 HflK subunits from different complexes could potentially interact with one or two FtsH hexamers, resulting in signal averaging that obscures FtsH density. Supporting this notion, lowpass filtering the map to 20 Å to attenuate high-frequency noise revealed a density at the cytoplasmic side of the complex, likely corresponding to the FtsH AAA and protease domain (Fig. S3C).

**Table 1.** Cryo-EM data collection, processing, model building, and validation statistics.

| name | Crosslinked HflK/C variant (HflK/C<sup>SS</sup>) | HflK/C<sup>WT</sup> solubilized in DDM from tobramycin-treated cells | HflK/C<sup>WT</sup> extracted in Carboxy-DIBMA from tobramycin-treated cells |
| --- | --- | --- | --- |
| PDB ID | 9PAW | NA |  |
| EMBD ID | EMD-71448 | EMD-71449 | EMD-71447 |
| Microscope | Titan Krios G3 | Titan Krios G3i | Titan Krios G3i |
| Camera | Falcon 4 | K3 detector (Gatan) | K3 detector (Gatan) |
| Magnification | 75,000 X | 130,000 X | 130,000 X |
| Voltage (kV) | 300 | 300 | 300 |
| Total electron dose (e<sup>-</sup>/Å<sup>2</sup>) | 52.68 e<sup>-</sup>/Å<sup>2</sup> | 48.02 e<sup>-</sup>/Å<sup>2</sup> | 46.86 e<sup>-</sup>/Å<sup>2</sup> |
| Defocus range (μm) | −1.2 to −2.4 μm | -0.5 to -1.75 μm | -1.3 μm |
| Pixel size (Å) | 0.8654 | 0.654 | 0.654 |
| Micrograph collected | 8,766 | 21,895 | 32,694 |
| Final particles | 299,919 | 58,175 | 24,196 |
| Symmetry | C<sub>1</sub> | C<sub>1</sub> | C<sub>1</sub> |
| Resolution (Å) GSFSC (0.143) | 2.9 | 9.1 | 11.5 |
| Unmasked resolution (Å) | 4.2 | 11 | 16 |
| Sphericity score (out of 1) | 0.85 | 0.90 | 0.80 |
| Model composition |  | NA |  |
| All atoms | 90,278 |  |  |
| Protein residues | 6433 |  |  |
| Ligands | 12 |  |  |
| Refinement |  |  |  |
| Map-model CC (mask) | 0.76 |  |  |
| RMSD bond lengths (Å) | 0.001 |  |  |
| RMSD bond angles (deg.) | 0.369 |  |  |
| Validation |  |  |  |
| MolProbity score | 1.18 |  |  |
| Clash score | 1.27 |  |  |
| C-beta outliers (%) | NA |  |  |
| Rotamer outliers (%) | 0.00 |  |  |
| Ramachandran favored (%) | 95.18 |  |  |
| Q-score (global/expected) | 0.41/0.53 |  |  |

**Figure 2.**
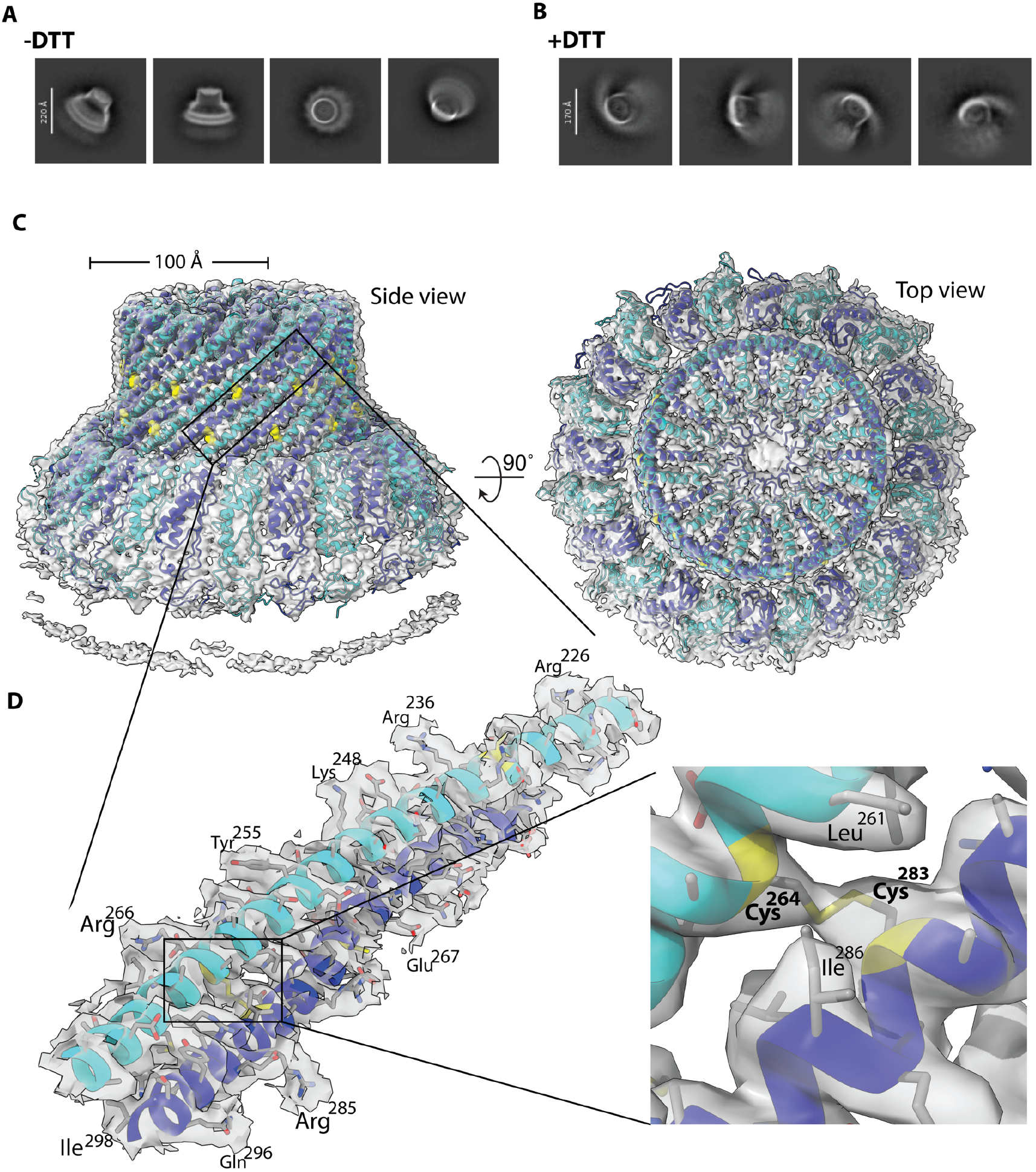
Cryo-EM structure of the DDM-solubilized, crosslinked FtsH•HflK/C^SS^ (HflK/C^SS^) complex. (**A**) 2D class averages of the HflK/C^**SS**^ variant under non-reducing conditions show the closed conformation of the assembly. **(B)** Under reducing conditions, HflK/C^**SS**^ adopts an open, nautilus-like conformation. (**C**) Unsharpened density map and cartoon models of HflK (blue) and HflC (cyan) viewed from the side and top, highlighting the closed HflK/C chamber. (**D**) Left panel: close-up view of the overlay between the atomic model and density map of the “hat” region of HflK (blue) and HflC (cyan), showing exemplary side chain density. Right panel: an exemplary density for the disulfide bond between HflK and HflC.

Comparison of HflK/C^**SS**^ structure with two previously published structures (PDB IDs: 9CZ2 and 7WI3)^5, 22^ revealed that the “hat” portion of the cage—where the disulfide bonds were introduced—was largely unaffected by cross-linking, except in the region where the opening originates in our earlier nautilus-like structure (Figs. 3A and 3B). However, a comparison between the two closed conformations—our current structure and a previously published one^22^—revealed a notable difference in the overall curvature of the HflK/C assembly (Figs. 3C–E). Since the ‘hat’ regions— where our disulfide bonds are located—were structurally similar, we speculate that the curvature observed in the earlier study (PDB ID: 7WI3) may have resulted from the use of the non-specific cross-linker glutaraldehyde during sample preparation^22^.

**Figure 3.**
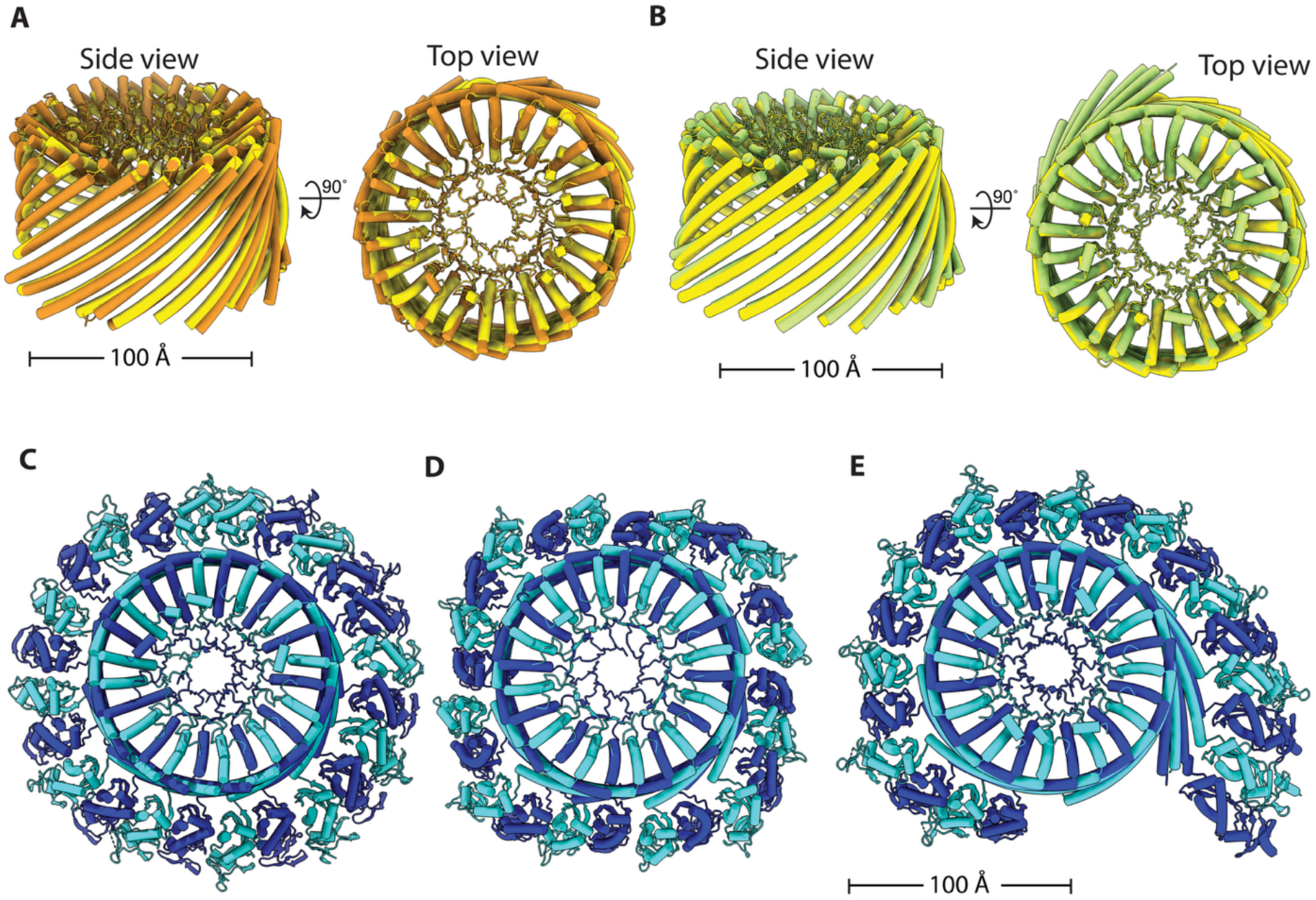
Comparison of the cross-linked HflK/C^SS^ closed conformation with previously determined closed and open structures. (**A**) Overlay of the “hat” region from HflK/C^**SS**^ (yellow) and the previously reported closed ‘cage’ structure (PDB ID: 7WI3, orange). The closed ‘cage’ structure was determined using both overexpression and glutar-aldehyde crosslinking. (**B**) Overlay of the “hat” region from HflK/C^**SS**^ (yellow) and the open, nautilus-like structure (PDB ID: 9CZ2, green). Only the HflK and HflC subunits located at the opening in the nautilus-like conformation exhibit conformational differences compared to the cross-linked HflK/C^**SS**^ structure. (**C–E**) Top-view comparisons of **(C)** the fully closed HflK/C^**SS**^ structure (**D**) the previously reported closed conformation (PDB ID: 7WI3); and (**E**) the nautilus-like open conformation of the native HflK/C complex (PDB ID: 9CZ2). All panels show cartoon representations with HflK in blue and HflC in cyan.

### Closed HflK/C conformation compromises bacterial stress response to aminoglycoside antibiotic stress

Having established a construct that exclusively favors the closed conformation, we next evaluated the impact of a closed HflK/C assembly on bacterial physiology under aminoglycoside stress. Although HflK/C is non-essential under normal conditions in *E. coli*, both FtsH and HflK/C are critical for recovery from aminoglycoside-induced stress, which compromises membrane integrity and leads to the accumulation of misfolded proteins requiring FtsH-mediated degradation^18-19^. To confirm previous studies showing that both *ftsH* and *hflK/C* are upregulated under such stress^19, 24^, we performed RNA-seq analysis comparing tobramycin-treated and untreated *E. coli* BL21 cells (Fig. S4). We observed upregulation of both *ftsH* and *hflK/C*, as well as known FtsH substrates such as *mgtA*, which is involved in regulating intracellular ionic strength^25-26^; *ibpA*, a small heat shock protein that buffers protein aggregation under heat or oxidative stress^27^; and other chaperones like ClpB and endopeptidases were also upregulated, suggesting an interplay between proteolysis and chaperone-mediated rescue (Fig. S4 and Source Data Table S1). To assess the functional relevance of the closed conformation of the HflK/C assembly, we performed plate-reader-based growth assays (Figs. 4 and S5) ^18^ in *BL21* Δ*hflK/C* cells complemented with either wild-type HflK/C (HflK/C^**WT**^), the cross-linked variant (HflK/C^**SS**^), or an empty vector (EV), all expressed from the same low-copy number plasmid used in our structural studies. In the absence of antibiotic stress, all strains grew similarly well at 37 °C (Fig. 4A), consistent with the non-essentiality of *hflK/C* under standard growth conditions^5, 20^. However, under tobramycin stress, only cells expressing HflK/C^**WT**^ recovered, while those expressing the cross-linked variant (HflK/C^**SS**^) or the empty vector (EV) showed significantly impaired growth (Fig. 4b).

**Figure 4.**
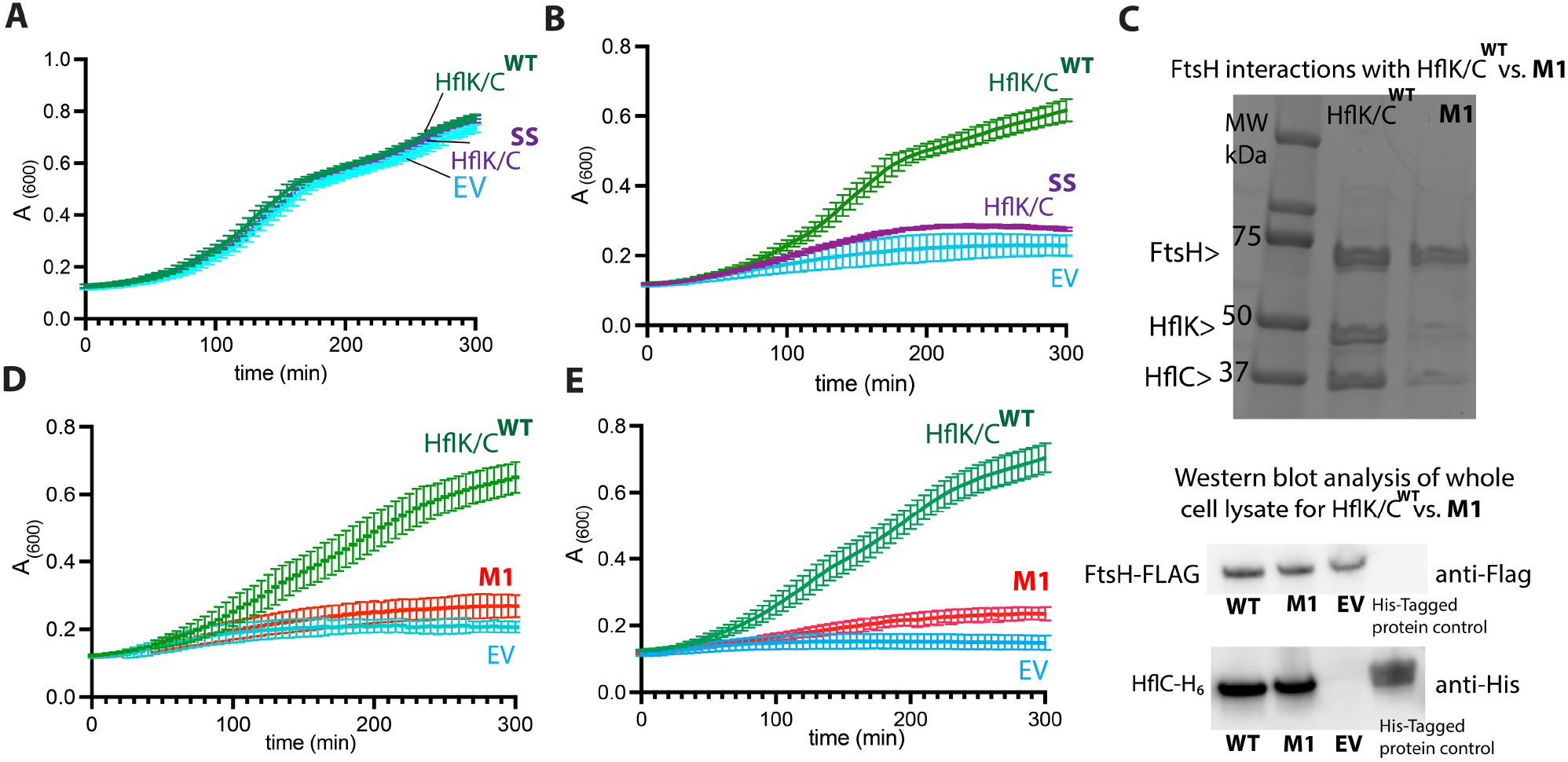
Effects of HflK/C variants on bacterial recovery under aminoglycoside stress. All assays were performed using *E. coli* BL21 Δ*hflK/C* cells expressing various HflK/C variants from a pPro24 plasmid under the control of a sodium propionate–inducible promoter at 37 °C in LB medium containing 50 µg/mL kanamycin, 35 µg/ml chloramphenicol, 1 mM sodium propionate unless otherwise noted. (**A**) Growth curves for cells expressing wild-type HflK/C (HflK/C^**WT**^), crosslinked HflK/C (HflK/C^**SS**^), and empty vector (EV). (**B**) Growth curves for cells expressing wild-type HflK/C (HflK/C^**WT**^), crosslinked HflK/C (HflK/C^**SS**^), and empty vector (EV) under tobramycin stress. (**C**) Top: Pulldown of the HflK/C complex solubilized in DDM from *E. coli* BL21 *ΔhflK/C* cells with chromosomally encoded FLAG-tagged FtsH and either wild-type or M1 (R141A, E142A, R185A) HflK, Bottom: Western blot analysis of the whole cell lysates shows a comparable expression of HflC between WT and M1 mutant. (**D–E**) Growth curves of tobramycin-stressed cells expressing: HflK/C^WT^, mutant **M1** [HflK(R141A, E142A, R185A)/HflC], and EV under 1mM sodium propionate (**D**); and 5 mM sodium propionate (**E**). Each growth assay was performed from a separate colony (n = 3 independent biological replicates), and data are presented as mean ± 1 SD.

### FtsH–HflK/C interactions are required for bacterial stress recovery

Previous studies have shown that FtsH associates with the HflK/C assembly exclusively through its interactions with HflK^14^, and that mutations in hydrophilic residues within the periplasmic domain of FtsH^22^ can significantly affect its interaction with the HflK/C complex. To investigate the functional role of this interaction in bacterial growth under antibiotic stress, we introduced mutations in HflK rather than in FtsH to avoid disrupting FtsH’s catalytic activity. Guided by our previous structures^5^, we substituted three hydrophilic residues in HflK (Arg141, Glu142, and Arg185) with alanine to disrupt potential contacts with FtsH (**M1** mutant; Fig. S6). The pulldown assay using Flag-tagged FtsH recovered HflK and HflC in the **M1** background at levels lower than wild type (Fig. 4C), suggesting that these residues are important for the FtsH–HflK/C interactions. To rule out the possibility that reduced recovery was due to impaired expression or stability of the HflK/C complex, we appended a Cterminal His-tag to HflC in both **WT** and **M1** backgrounds. As previously reported^28^, HflC is unstable in the absence of HflK, and thus serves as a readout for HflK/C complex formation. Immunoblot analysis revealed similar levels of HflC in **WT** and **M1** strains, confirming that the **M1** mutations do not compromise HflK/C level. Functionally, cells expressing the **M1** variant from a propionate-titratable promoter at both low (1 mM) and high (5 mM) sodium propionate concentrations showed impaired recovery under tobramycin stress compared to those expressing HflK/C^**WT**^ (Figs. 4D, 4E, and S7), supporting a critical role for the FtsH–HflK/C interaction in promoting stress adaptation.

### The HflK/C^WT^ adopts a new conformation under antibiotic stress

Since the fully closed HflK/C assembly and mutations that disrupt FtsH–HflK/C interactions led to reduced bacterial growth under antibiotic stress, we conclude that the flexible, open conformation of HflK/C and its interaction with FtsH are essential for regulating FtsH proteolysis during tobramycin-induced stress. To investigate how the HflK/C complex structurally aids bacterial growth under antibiotic stress, we exposed bacteria to tobramycin and subsequently isolated the endogenous FtsH complex using either the micelle-forming detergent DDM or, in a detergent-free approach, the nanodisc-forming polymer Carboxy-Diisobutylene-Maleic Acid (Carboxy-DIBMA), which we have previously employed for FtsH•HflK/C structural studies^5^. Surprisingly, structural analysis of the purified complexes (Figs. S8–11; Table 1) revealed a secondary opening in the HflK/C assembly, located opposite the previously identified primary aperture and positioned near the second FtsH hexamer (Fig. 5). The width of this opening (∼30–50 Å) is narrower than that of the initial aperture (∼70-100 Å) yet still sufficiently wide to permit the diffusion of small membrane proteins into the assembly. This secondary opening was observed in both DDM micelles and Carboxy-DIBMA nanodiscs; however, only the latter additionally revealed the presence and orientation of the adjacent lipid bilayer. In addition to the density corresponding to the periplasmic side of FtsH, we also observed extra density surrounding FtsH, which we suspected to represent substrate. However, the resolution of these maps (∼9–11 Å) was insufficient to draw definitive conclusions. To evaluate the structural heterogeneity of the complex, we performed cryoDRGN analysis^29-30^, which revealed variability in both the aperture widths and the positioning of FtsH within the assembly (Movie 1).

**Figure 5.**
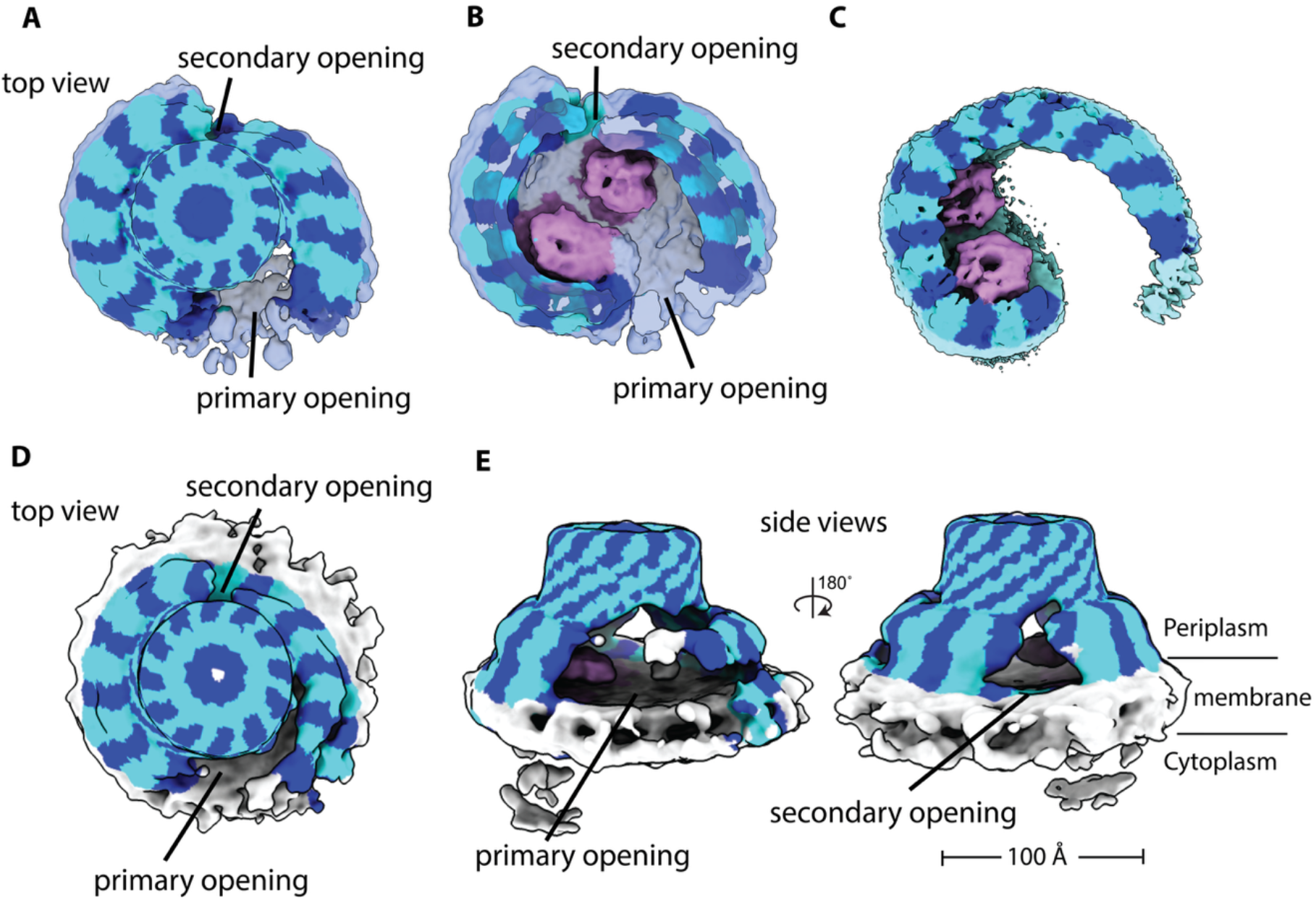
Cryo-EM maps of the native HflK/C complex from cells treated with tobramycin. The maps are color-coded using an atomic model: HflK in blue, HflC in cyan, and FtsH in pink. (**A, B**) Cryo-EM reconstructions of DDM-solubilized HflK/C complexes showing a top view and a sliced top view, respectively. Two openings are observed near the positions where FtsH hexamers are embedded within the HflK/C assembly. (**C**) Previously determined cryo-EM map of the nautilus-like FtsH•HflK/C complex (EMD-46057). (**D**) Cryo-EM maps of HflK/C complexes extracted into Carboxy-DIBMA nanodiscs, showing the secondary opening toward the lipid bilayer, as seen from both top and side views.

## DISCUSSION

The HflK/C membrane assembly, a member of the SPFH protein family, and the AAA+ protease FtsH together play a key role in bacterial stress responses, particularly in mediating resistance to aminoglycosides^18-19, 31^. Despite their importance, the mechanisms by which HflK/C support bacterial recovery by modulating FtsH activity remain poorly understood^19, 23^. Additionally, recent structures of HflK/C in two distinct conformations—an asymmetric, open, nautilus-like structure^5^ and a symmetric, closed, cage-like structure^22-23^—have raised new questions about the biologically active conformation of HflK/C and its role in regulating the proteolytic activity of FtsH. By engineering disulfide bonds into the ‘hat’ region of HflK/C, we were able to stabilize the closed conformation (Fig. 1 and Fig. 2). The cryo-EM structure revealed that the closed conformation observed in this crosslinked variant of HflK/C resembles the closed conformations of other SPFH family structures^32-33^ but differs from the previously reported closed FtsH•HflK/C complex^22^(Figs. 3C–E). We propose that this discrepancy may stem from the use of a non-specific crosslinker in the previous study^22^, which could have altered the native arrangement of the SPFH domains and resulted in a cubic cage structure rather than a rounded one observed here (Fig. 3). Our phenotypic results showed that HflK/C in the closed conformation significantly impairs bacterial growth under aminoglycoside stress (Figs. 4a and 4b), suggesting that this conformation represents an inactive state of the HflK/C assembly during membrane stress.

However, our phenotypic assay did not reveal any growth defect under normal conditions for the fully closed variant of HflK/C (Fig. 4a), which is consistent with the fact that *hflK* and *hflC* are nonessential under standard growth conditions^5^. This suggests that the closed conformation still permits FtsH to degrade essential cytoplasmic substrates, such as LpxC^3^, while damaged or unnecessary membrane proteins may be eliminated in dividing cells either through dilution during cell division or via alternative proteolytic pathways, including endopeptidases or HflK/C-independent degradation by FtsH^34^. In contrast, the open conformation may be specifically required during conditions that result in the accumulation of excessive membrane-associated substrates.

While the question of how the HflK/C assembly contributes to substrate recruitment by FtsH remains open, our mutational analysis of HflK suggests that specific interactions between FtsH and HflK are critical for regulating FtsH proteolysis to support bacterial growth under aminoglycoside stress (Fig. 4C–4E). These interactions may facilitate the spatial organization of FtsH and its substrates within the HflK/C complex, potentially enhancing substrate recognition by the AAA domain through an increased local concentration of substrates around two FtsH hexamers. Consistent with this model, the cryo-EM structure of the native FtsH•HflK/C complex isolated from aminoglycoside-treated cells revealed an unprecedented opening near the second FtsH (Fig. 5). This aperture likely facilitates access to both FtsHs under proteotoxic stress, allowing excess unfolded protein substrates to enter the microdomain from multiple directions. Whether this opening results from post-translational modifications, proteolytic cleavage by endopeptidases, or arises passively in response to oxidative membrane damage^19^ remains to be determined. This result, together with our previous proteomic analysis^5^, supports a model in which dynamic opening of HflK/C fine-tunes FtsH-mediated proteolysis, particularly during aminoglycoside-induced membrane stress, when misfolded or damaged membrane proteins accumulate and must be rapidly cleared by the FtsH to restore cellular homeostasis^19, 35^.

Interestingly, other members of the SPFH family have also been implicated in stress recovery mechanisms across diverse organisms^21, 36-39^. For instance, stomatin contributes to mitigating lipopolysaccharide-induced oxidative stress and inflammation in the mouse lung^37^; flotillins stabilize unfolded proteins in methicillin-resistant *Staphylococcus* under cellular stress^36^; and prohibitin serves as a biomarker of oxidative stress in ocular tissues^39-40^. Given the shared structural and functional features, could the stress-induced structural changes observed in HflK/C represent a conserved property among SPFH family members across species?

Recent cryo-EM structures of other SPFH proteins have predominantly revealed closed assemblies^23, 41-43^. However, the predominance of closed conformations observed in other SPFH proteins may, in part, reflect experimental factors such as protein overexpression^22-23, 41^, the imposition of high symmetry during 3D cryo-EM reconstruction (often applied to a subset of symmetrical particles following extensive 3D classifications)^23, 41-42^, or using non-specific cross-linkers^22^—all of which could bias structural determination toward a closed state. Another possibility is that the conformational flexibility observed in HflK/C may represent a unique feature of SPFH complexes that interact with AAA+ proteases. Supporting this notion, a recent preprint using *in situ* cryo-ET of mitochondria under depolarizing conditions showed that prohibitins—the mitochondrial homologs of HflK/C—can adopt both an asymmetric, open, nautilus-like conformation and a symmetric, closed arrangement^44^ similar to our closed HflK/C structure presented here, although m-AAA was not observed in those assemblies. Our use of targeted disulfide cross-linking to selectively stabilize the closed HflK/C conformation and demonstrate its physiological impact provides an approach that could be extended to other SPFH family members to probe the biological relevance of the closed state.

Overall, our study provides novel insights into how HflK/C conformational dynamics contribute to bacterial recovery from aminoglycoside-induced stress and reveals how this massive membrane complex structurally adapts to proteotoxic conditions. Our study may provide the groundwork for understanding the functional mechanisms of SPFH family proteins in both bacteria and eukaryotes, as well as their roles in cellular stress response pathways.

## MATERIALS AND METHODS

### Plasmid construction

Plasmids were generated by PCR amplification of the region spanning 170bp upstream of *hflK* to 51bp downstream of *hflC* from the *E. coli* BL21 chromosome and cloning into a plasmid containing a propionate-inducible promoter^45^. Additional mutations corresponding to the HflK/C^**SS**^ variant (A270C and A283C in HflK; A233C and A264C in HflC) and the **M1** variant of HflK (R141A, E142A, R185A) were introduced using synthetic gene blocks (GenScript Inc.) and assembled by Gibson assembly. A C-terminally His-tagged version of *hflC* was generated by Q5 site-directed mutagenesis following the manufacturer’s protocol.

### FtsH•HflK/C purification

Previously constructed strains containing a C-terminal FLAG-tagged variant of FtsH^5^, introduced at the endogenous *ftsH* locus in *E. coli* BL21(DE3) or BL21(DE3) Δ*hflK/C*, were used for all pull-down experiments. A single bacterial colony was inoculated into 50 mL of LB medium containing 50 µg/mL kanamycin and 35 µg/mL chloramphenicol and grown overnight at 37°C with shaking. Overnight cultures were inoculated into 4 L of fresh medium (50% LB:50% YT) and incubated at 37 °C with shaking at 220 rpm for 24 h. Cells were harvested by centrifugation at 4,000 rpm for 25 min at 4 °C using an H-1200 rotor. The resulting pellet was resuspended in Buffer A (100 mM KCl, 50 mM Tris-HCl, pH 8.0, 10% glycerol, 5 mM MgCl_2_, 100 µM ZnCl_2_) and stored at –80 °C until further use. For lysis, frozen cells were thawed on ice and disrupted by sonication (30% amplitude, 10 s on/30 s off cycles, for a total of 3 min). Cell debris was removed by centrifugation at 14,000 rpm for 20 min at 4 °C using a JL20 rotor. The supernatant was ultracentrifuged at 30,000 rpm for 1 h at 4 °C in a 45 Ti rotor to isolate the membrane fraction. The resulting membrane pellet was resuspended in Buffer A and homogenized using a Dounce homogenizer. A 2% DDM stock solution was added to achieve a final DDM concentration of 1%, and the mixture was incubated at 4 °C for 2 h to allow membrane protein solubilization. Samples were then ultracentrifuged at 30,000 rpm for 1 h at 4 °C, and the clarified supernatants were collected and kept on ice. The samples were incubated with pre-washed M2-FLAG resin (Millipore Sigma, Cat. #A2220). The solubilized protein solution was added to the prepared resin and incubated at 4°C for 2 h with gentle rotation. After incubation, the resin was centrifuged at 400 *g* for 5 min, loaded into a gravity column, and washed with 2 mL of Buffer A (in 200 µL aliquots) containing 0.03% DDM. A second wash was performed using Buffer B (400 mM NaCl, 50 mM Tris-HCl, 5 mM MgCl_2_, 10% glycerol, and 0.03% DDM). The samples were eluted using a 0.25 mg/mL Flag peptide (APEXBio) in Buffer B (100 mM KCl, 50 mM Tris-HCl pH 8.0, 5% glycerol, 5 mM MgCl_2_, 100 µM ZnCl_2_, 0.03% DDM). The pooled elution fractions were concentrated using a 100 kDa cutoff Centricon filter (Millipore Inc.). The same protocol was used for GDN sample preparation, except 2% GDN was used for membrane solubilization and 0.03% GDN was employed for the wash and elution steps. For Carboxy-DIBMA and DDM extractions from tobramycin-treated cells, the previously described protocol was followed^5^. The only difference was that cells were treated with 2.5 µg/mL tobramycin overnight before complex extraction and purification.

### Phenotypic assay

Cellular assays were performed using previously constructed *E. coli* BL21(DE3) Δ*hflK/C* cells carrying a chromosomally encoded C-terminal FLAG tag at the *ftsH* locus^5^. All plasmids, or an empty vector control, were transformed into BL21(DE3) Δ*hflK/C* cells and assayed as described below. For growth assays, overnight cultures were grown at 37 °C and then diluted to an A_600_ of ∼0.1 in LB medium containing chloramphenicol (35 μg/mL) and 1 mM sodium propionate. Aliquots (200 μL) of bacterial suspensions containing different *hflK/C* variants or an empty vector were transferred to a 96-well clear flat-bottom microplate (Corning Inc.). Absorbance at 600 nm (A_600_) was measured every 5 min using a SpectraMax M2 plate reader.

### Pull-down assay

Overnight cultures of cells expressing either wildtype or mutant constructs (using the same plasmids employed for structural and phenotypic assays) were inoculated into 1 L of fresh medium consisting of a 1:1 mixture of LB and YT. Cultures were incubated at 37 °C with shaking for 24 h. Cells were harvested by centrifugation at 4,000 rpm for 30 min at 4 °C, and cell pellets were resuspended in 30 mL of buffer A and stored at – 80°C. Upon thawing, cells were lysed by sonication (30% amplitude, 10 seconds on / 30 seconds off, total sonication time of 3 minutes). The lysate was centrifuged at 14,000 *g* for 30 min using a JA20 rotor to remove cell debris. The cleared supernatant was then ultracentrifuged at 30,000 rpm for 1 h in a Ti-45 rotor to isolate the membrane fraction. The membrane pellet was resuspended in 6 mL of Buffer A and homogenized with 5–6 strokes of a Dounce homogenizer. An equal volume of 2% DDM was added to reach a final concentration of 1% DDM, and the suspension was incubated at 4°C for 2 h with gentle mixing to solubilize membrane proteins. Following solubilization, the sample was ultracentrifuged again at 30,000 rpm using 45Ti rotor for 1 h, and the supernatant containing solubilized proteins was collected and kept on ice. For affinity purification, 250 µL of M2-FLAG resin was washed with 2 mL Buffer A containing 0.03% DDM and pelleted by centrifugation at 400 *g* for 5 min. The solubilized protein solution was incubated with the prewashed resin (with buffer A) at 4°C for 2 h with gentle mixing. After incubation, the resin was pelleted by centrifugation at 400 *g*, transferred to a gravity-flow column, and washed sequentially with 2 mL Buffer A (+0.03% DDM) and 2 mL Buffer B (+ 0.03% DDM). Elution was performed using 0.25 mg/ml Flag peptide (APEXBio) diluted in Buffer C (100 mM KCl, 50 mM Tris-HCl pH 8.0, 5% glycerol, 5 mM MgCl_2_, 100 µM ZnCl_2_, 3 mM β-mercaptoethanol, 0.03% DDM). Eluates were pooled and concentrated using a 100 kDa cutoff centrifugal filter unit (centrifuged at 6,000 rpm at 4°C). For SDS-PAGE analysis, 4 µL of each sample fraction was mixed with 10 µL of 4× Laemmli buffer, boiled for 5 min, and 6 µL of the mixture was loaded onto the 4–20% Mini-PROTEAN TGX Precast Protein Gels.

### Cryo-EM sample preparation

For the cross-linked FtsH•HflK/C (FtsH•HflK/C^**SS**^) structure, samples were prepared by applying 2.5 μL of ∼0.3 mg/mL of FtsH.HflK/C (crosslinked) complex on 200 mesh Quantifoil 2 nm carbon 2/1 copper grids. The grids were glow-discharged using a GloQube Plus (MiTeGen) at 15 mA for 20 s. Sample loaded grids were blotted for 4s with a blot force of +4 at 6°C and 100 % relative humidity using a FEI Vitrobot Mark IV instrument (Thermo Scientific).

For the DDM-solubilized FtsH•HflK/C complex extracted from the tobramycin treated cells, ∼0.7 mg/mL HflK/C sample was used for grid preparation. 2-nm carbon-supported 200-mesh Quantifoil 2/1 copper grids, which had been glow-discharged for 20 s in an easiGlow glow discharger (Pelco) at 15 mA, were utilized. For the Carboxy-DIBMA-extracted sample from the tobramycin treated cells, a concentration of ∼0.3 mg/mL of the FtsH•HflK/C complex was applied to 2-nm carbon-supported 200-mesh Quantifoil 2/1 copper grids, also glow-discharged for 20 s in an easiGlow glow discharger (Pelco) at 15 mA, and the sample was vitrified as above.

### Cryo-EM data collection

For the cross-linked FtsH•HflK/C (FtsH•HflK/C^**SS**^) complex structure, 8,766 movies were collected with EPU on a Titan Krios G3 using multiple images per hole with an acceleration voltage of 300 kV and magnification of 75k, detected on a Falcon 4 detector for an effective pixel size of 0.8654 Å. Movies were collected as 50 frames with a defocus range from −1.2 to −2.4µm and a total exposure per specimen of 52.68 e^-^/Å^2^.

For the DDM-solubilized FtsH•HflK/C complex extracted from the tobramycin treated cells, 21,895 movies were collected with EPU using aberration-free image shift (AFIS) and hole-clustering method on a Titan Krios G3i with an acceleration voltage of 300 kV and magnification of 130,000×, detected in super-resolution mode on a Gatan K3 detector for an effective pixel size of 0.654 Å (binned by 2). Movies were collected as 40 frames with a defocus range from −0.5 to −1.75µm and a total exposure per specimen of 48.02 e^-^/Å^2^.

For the Carboxy-DIBMA-extracted FtsH•HflK/C sample from tobramycin-treated cells, 32,907 movies were collected as 40-frame, with a defocus range of −1.3 µm and a total electron dose of 46.40 e^−^/Å^2^ per specimen. Data acquisition was performed using AFIS hole clustering at a 25° stage tilt on a Titan Krios G3i operated at 300 kV acceleration voltage and 130,000× magnification. Images were recorded in super-resolution mode on a Gatan K3 detector, yielding an effective pixel size of 0.654 Å (binned by 2).

### Cryo-EM pre-processing and particle picking

For the DDM-solubilized FtsH•HflK/C^**SS**^ complex, data processing was performed using cryoSPARC (v4.6) and default parameters unless noted. Raw movies (8,766) were pre-processed using “Patch motion correction”, and “Patch CTF estimation”. Particles (429, 520) were picked using the “Blob-picker” tool applied to 3000 micrographs selected at random. Particles were extracted (box size 640 px, Fourier-cropped to 360 px) and classified using the ‘2D classification’ utility. The entire set of micrographs were then picked with the “Template picker” tool, using classes from 2D classification. 1,866,826 particles were extracted (box size 600 px, Fourier-cropped to 256 px) and subjected to multiple rounds of 2D classification, resulting in the selection of 309,350 particles as a preliminary stack.

For the DDM-solubilized FtsH•HflK/C complex extracted from the tobramycin treated cells, data processing was performed using cryoSPARC (v4.5)^46^ and default parameters unless noted. Raw movies (21,895) were pre-processed using “Patch motion correction”, and “Patch CTF estimation”. Particles (1,089,115) were picked using the “Blob-picker” tool applied to 3000 micrographs selected at random. After two rounds of 2D classification, 38 2D classes were selected as a template. Particles were picked using “Template picker” and were extracted (box size 900 px, Fourier-cropped to 220 px) and classified using the ‘Ab-initio’ utility (3 classes). 382,188 particles were selected and subjected to multiple rounds of 2D classification, resulting in the selection of 146,192 particles as a preliminary stack. For the Carboxy-DIBMA-extracted FtsH•HflK/C sample from tobramycin-treated cells, 32,694 movies were pre-processed using “Patch motion correction” and “Patch CTF estimation.” A total of 2,462,066 particles were initially picked using the “Blob picker.” After multiple rounds of 2D classification, three 2D classes were selected as templates. Particles were then repicked using the “Template picker” and extracted (box size: 900 px, Fourier-cropped to 128 px). After four additional rounds of 2D classification, 43,479 particles were selected as the preliminary stack.

### Ab initio reconstruction and global refinement

For the FtsH•HflK/C^**SS**^ complex solubilized in DDM, “Ab-initio reconstruction” using 1 class was performed, resulting in a model and the particle stack (300,000 particles) was selected for subsequent “non uniform refinement”. Particles were then re-extracted using 512 px box size down sampled to 384 px. Extracted particles were then run through non uniform refinement with C_4_ symmetry applied followed by another round of nonuniform refinement with C_1_ symmetry. The particles were then run through local refinement with C_1_ symmetry. A mask was then created in ChimeraX and local refinement was performed using this mask. After applying global and local CTF refinement followed by local refinement with C_1_ symmetry, the final map achieved at GSFSC resolution of ∼2.93 Å after FSC mask auto-tightening. Model building was carried out using ChimeraX^47^ (v1.9), Coot^27^ (v0.9.4), and Phenix^48^ (v1.21).

For the DDM-solubilized FtsH•HflK/C complex extracted from tobramycin-treated cells, *ab initio* reconstruction with three classes was performed. One particle class (58,175 particles) was selected for subsequent non-uniform refinement followed by local refinement. The final map achieved at GSFSC resolution of 9.1 Å after FSC mask auto-tightening. For the Carboxy-DIBMA-extracted FtsH•HflK/C sample from tobramycin-treated cells, *ab initio* reconstruction was performed using two classes. The class showing the FtsH•HflK/C complex was selected for non-uniform refinement leading to the final map achieved a GSFSC resolution of ∼11.5 Å after FSC mask auto-tightening.

### Western blot analysis

Western blot was performed by pelleting 1 mL of cells expressing either the HflK/C^**WT**^ or **M1** mutant at an A_600_ of 3.0 and 3.2 via centrifugation at 17000 *g* for 3 min. The supernatant was discarded, and cell pellets were resuspended in Laemmli buffer, supplemented with 1× EDTA-free protease inhibitor cocktail (Complete, Roche). Samples were boiled for 5 min and analyzed by SDS-PAGE. The bands were transferred onto a pre-activated PVDF membrane (Cat. #1704156) using the Trans-Blot Turbo transfer system (Bio-Rad). The membrane was incubated with 6x-His Tag Monoclonal Antibody (Cat. #MA1-21315; Thermo Fisher Scientific Inc.;1:5000 dilution), followed by goat anti-mouse IgG secondary antibody (Cat. #152-3920; Thermo Fisher Scientific Inc; 1:10,000 dilution). Signal was detected using Clarity Western ECL substrate (Bio-Rad, Cat. #170-5060) and visualized with a Bio-Rad imaging system.

### RNA-seq sample preparation and analysis

*E. coli* BL21 cells expressing FLAG-tagged wild-type FtsH were grown to an A_600_ of 0.5 in the presence or absence of 2.5 µg/mL tobramycin. Total RNA was extracted using the RNeasy Plus Mini Kit (Qiagen, cat. no. 74134) and processed at the Genome Technology Access Center, McDonnell Genome Institute, Washington University School of Medicine. RNA-seq libraries were prepared following the manufacturer’s protocol, indexed, pooled, and sequenced on an Illumina NovaSeq X Plus. Base calling and demultiplexing were performed using Illumina DRAGEN and BCLconvert (v4.2.4).

Reads were aligned to the primary genome assembly using STAR (v2.7.11b)^49^. Gene-level counts were obtained using Subread:featureCounts (v2.0.8)^50^, and transcript-level quantification of known Ensemble isoforms was performed with Salmon (v1.10.0)^51^. Sequencing quality metrics— including total aligned reads, uniquely mapped reads, and detected features—were assessed. Ribosomal RNA content, splice junction saturation, and read distribution across gene models were evaluated using RSeQC (v5.0.4)^52^.

Gene counts were imported into the EdgeR package (Bioconductor v4.4.0)^53^ and normalized using the trimmed mean of M-values (TMM) method. Genes corresponding to ribosomal RNAs and those not expressed at ≥1 count-per-million in at least three samples were excluded. The resulting TMM-normalized counts were analyzed using Limma^54^. Weighted likelihoods, accounting for the mean-variance relationship across genes, were calculated with the voomWithQuality Weights function^55^ used to fit a linear model. Model performance was assessed by plotting the residual standard deviation versus average log-counts, with a robustly fitted trend line. Differential expression was evaluated using empirical Bayes moderation, and significance was determined by applying Benjamini-Hochberg correction (FDR ≤ 0.05).

### CryoDRGN analysis

To investigate structural heterogeneity, we analyzed all 41,259 particles from the carboxy-DIBMA dataset using CryoDRGN v2.3.0^29-30, 56^. Particles were downsampled to a box size of 128 pixels (∼5.1 Å/pixel) and used to train an eight-dimensional latent-variable model with 1024×3 encoder and decoder architectures. Particle poses and CTF parameters for CryoDRGN training were obtained from the non-uniform refinement. After 50 epochs of training, 100 volumes were sampled from the k-means cluster centers of the latent embeddings. Visual inspection of the resulting volumes, guided by our atomic model, revealed structural variability in the opening length, membrane span, and orientation of FtsH within the assembly and relative to the lipid bilayer.

## Author Contributions

N.I. and A.G. expressed and purified proteins. N.I. and A.G. prepared samples for EM imaging and collected data. N.I. and A.G. processed EM data and performed reconstruction and refinement. N.I. conducted all phenotypic growth assays. SK provided a panel of DIBMA polymers and guidance in performing detergent-free extractions. A.G. built and refined the models and wrote the first draft. All authors contributed to writing and editing the manuscript. A.G. supervised the project.

## ACKNOWLEDGEMENT

This work was supported by startup funds from the Department of Biochemistry and Molecular Biophysics at Washington University School of Medicine and NIH grant R35-GM141517. We thank the Genome Technology Access Center at the McDonnell Genome Institute at Washington University School of Medicine for help with genomic analysis. The Center is partially supported by NCI Cancer Center Support Grant #P30 CA91842 to the Siteman Cancer Center from the National Center for Research Resources (NCRR), a component of the National Institutes of Health (NIH), and NIH Roadmap for Medical Research. This publication is solely the responsibility of the authors and does not necessarily represent the official view of NCRR or NIH. Cryo-EM data were collected at the Washington University in St. Louis Center for Cellular Imaging (WUCCI) and Cryogenic Electron Microscopy Facility at MIT.nano, which was a gift from the Arnold and Mabel Beckman Foundation. We thank all members of the Ghanbarpour and Keller labs as well as Bob Sauer and Maria Carreira for their helpful advice and discussions. We thank SBGrid^57^ for providing the structural biology software packages used in this study.

