## Supplementary material for "Structural Plasticity of the Membrane-Bound Protein Degradation Assembly Supports Bacterial Adaptation to Stress": SI: Supporting Information-tobo-bioRX-red.pdf

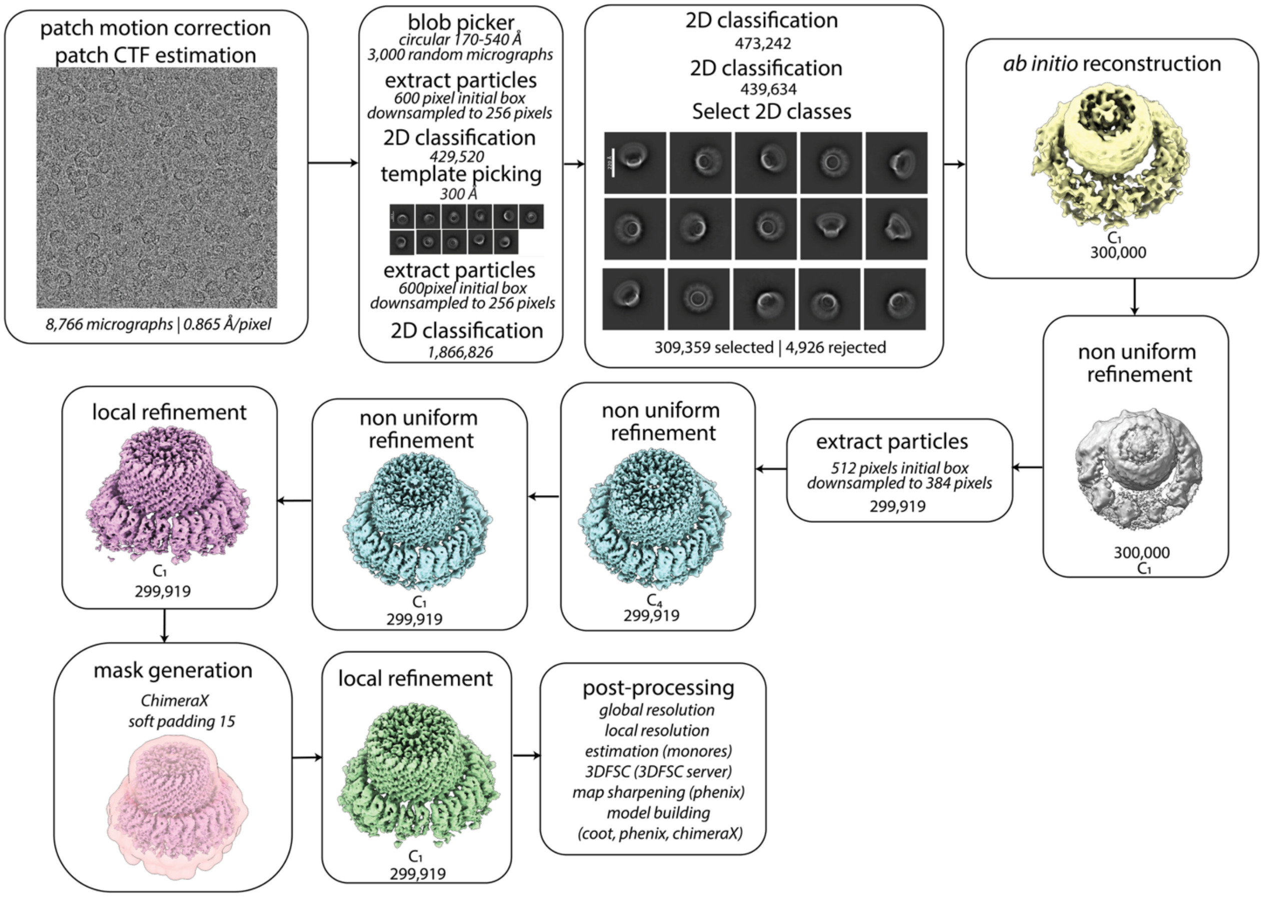


**Supplementary Figure 1. CryoSPARC processing workflow for cross-linked HflK/C$^{SS}$ complex.** Job names, job details, and non-default parameters (italicized) are noted in each box.

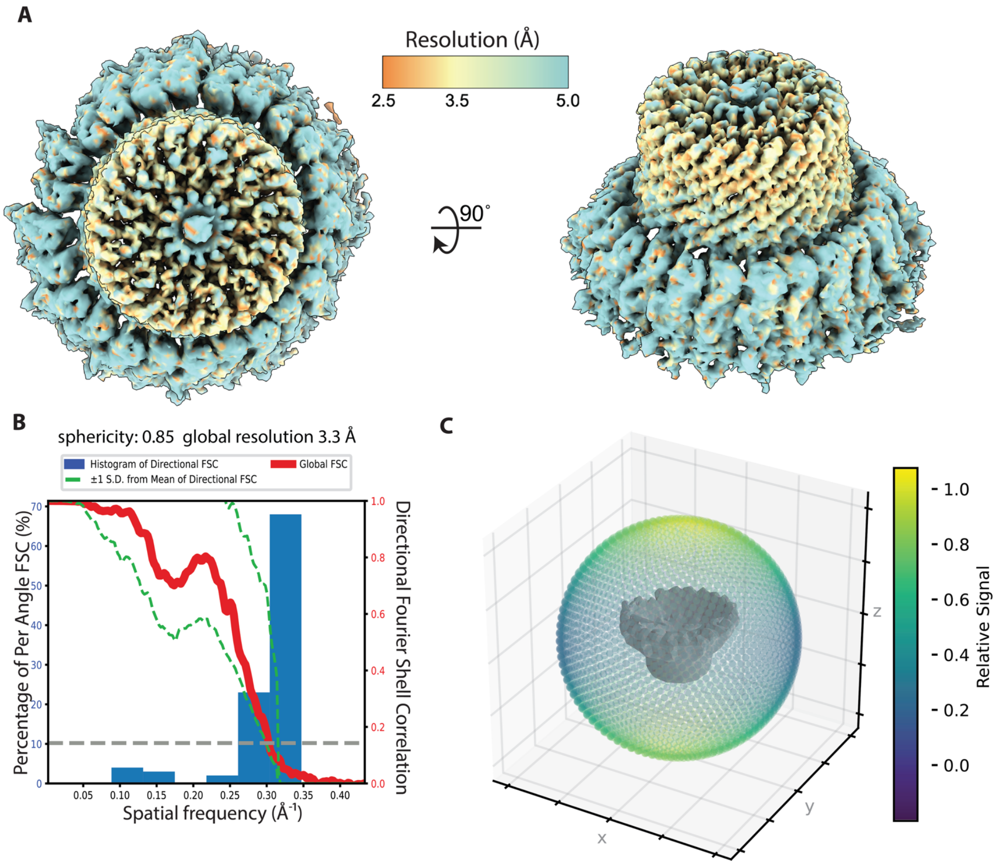


**Supplementary Figure 2. Estimates of resolution and angular sampling: DDM-solubilized FtsH•HflK/C$^{SS}$ complex.** **(A)** Maps colored by local resolution as estimated by the cryoSPARC implementation of monoRes. **(B)** Global resolution and directional resolution calculated by 3DFSC server (https://3dfsc.salk.edu). **(C)** Projection angle distribution estimated by cryoSPARC.

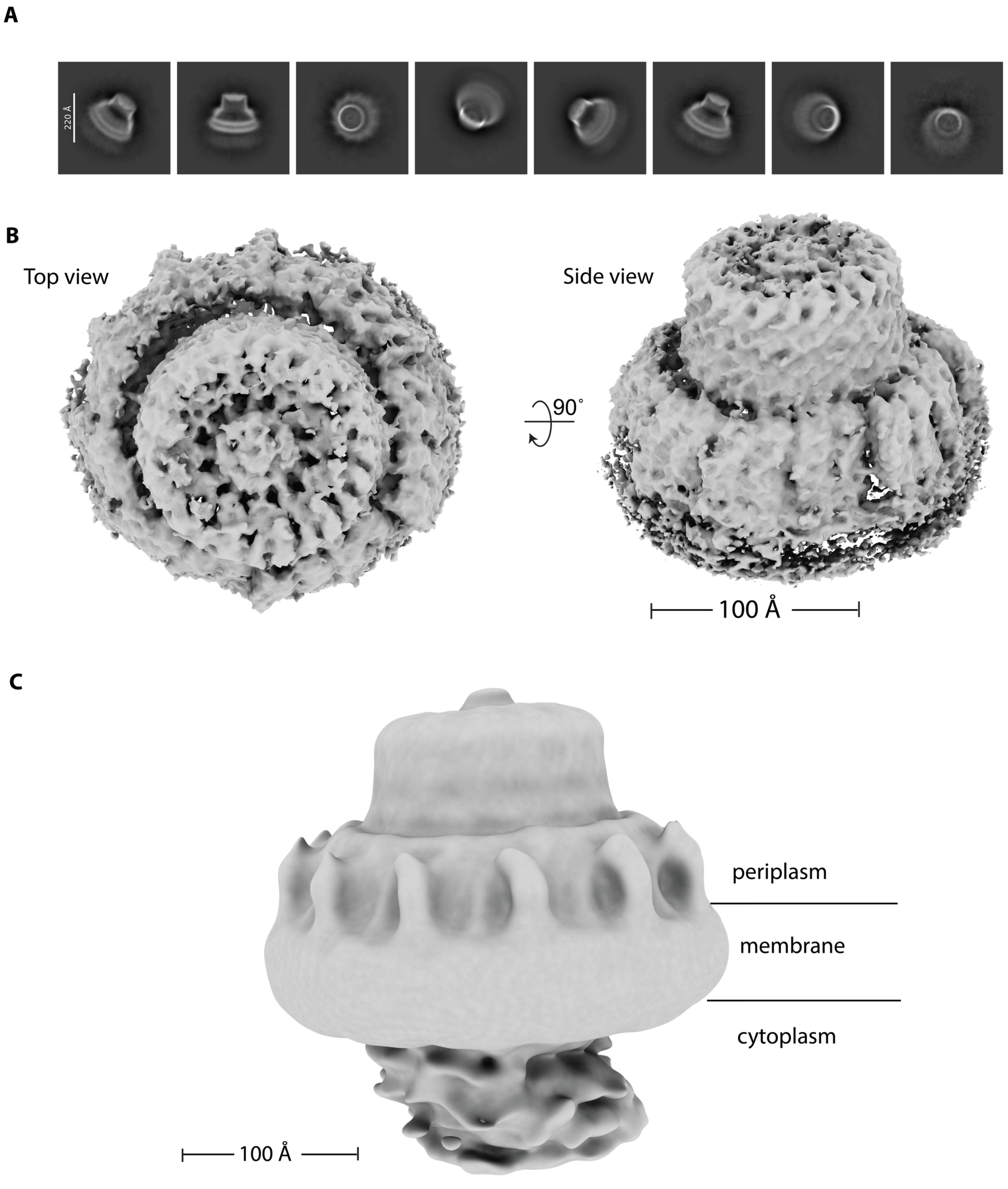


**Supplementary Figure 3. Cryo-EM structure of the crosslinked, GDN-solubilized FtsH–HflK/C (HflK/C$^{SS}$) complex and low-pass-filtered map of the DDM-solubilized HflK/C$^{SS}$ complex.** (**A**) Representative 2D class averages of the crosslinked complex. (**B**) Cryo-EM map of the final 3D reconstruction, resolved at global GS-FSC resolution: 4.14 Å. (**C**) Cryo-EM map of the DDM-solubilized HflK/C$^{SS}$ complex, low-pass filtered to 20 Å to enhance visibility of low-resolution features.

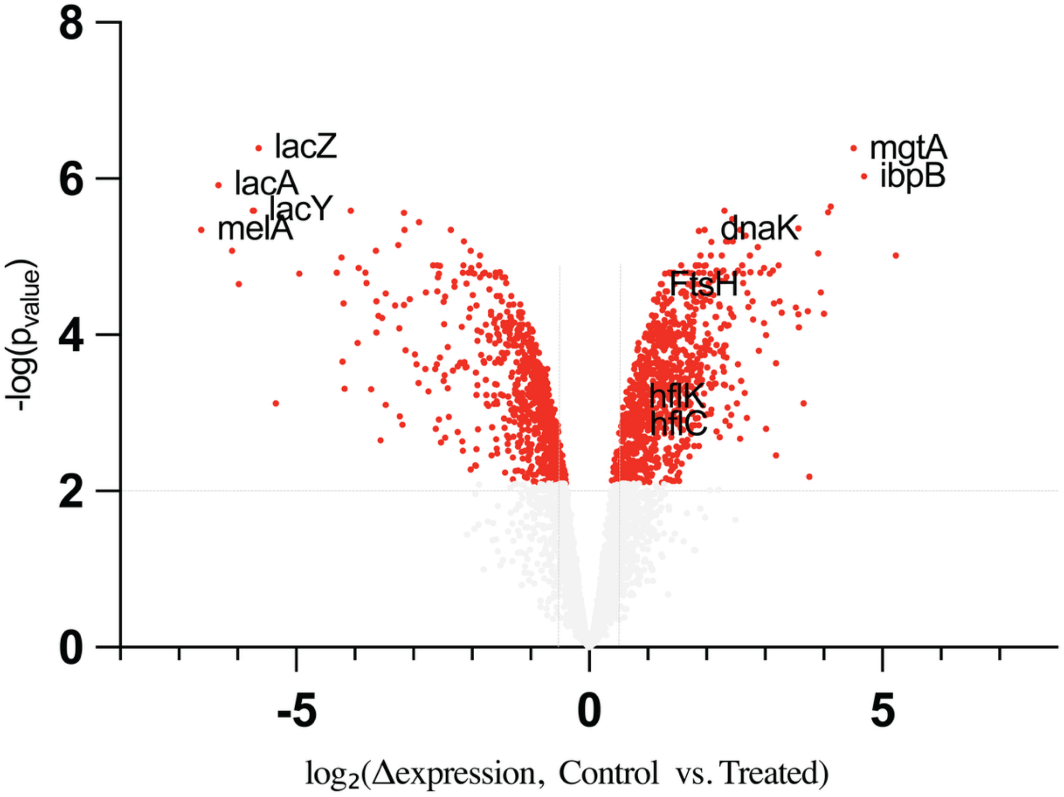


**Supplementary Figure 4. Volcano plot of RNA-seq data comparing control and tobramycin-treated *E. coli* BL21 cells.** Each point represents a gene, plotted by $\log_2$ fold change (x-axis) and statistical significance ($-\log P$-value, *y*-axis). Genes with $-\log P > 2$ and $\log_2$ $\Delta$expression $> 0.5$ are considered significantly upregulated (red). Notably upregulated genes include *ftsH*, *hflK*, *hflC*, *mgtA*, *ibpB*, and *dnaK*, a chaperone known to interact with FtsH. Downregulated genes include *lacZ*, *lacA*, *lacY*, and *melA*.

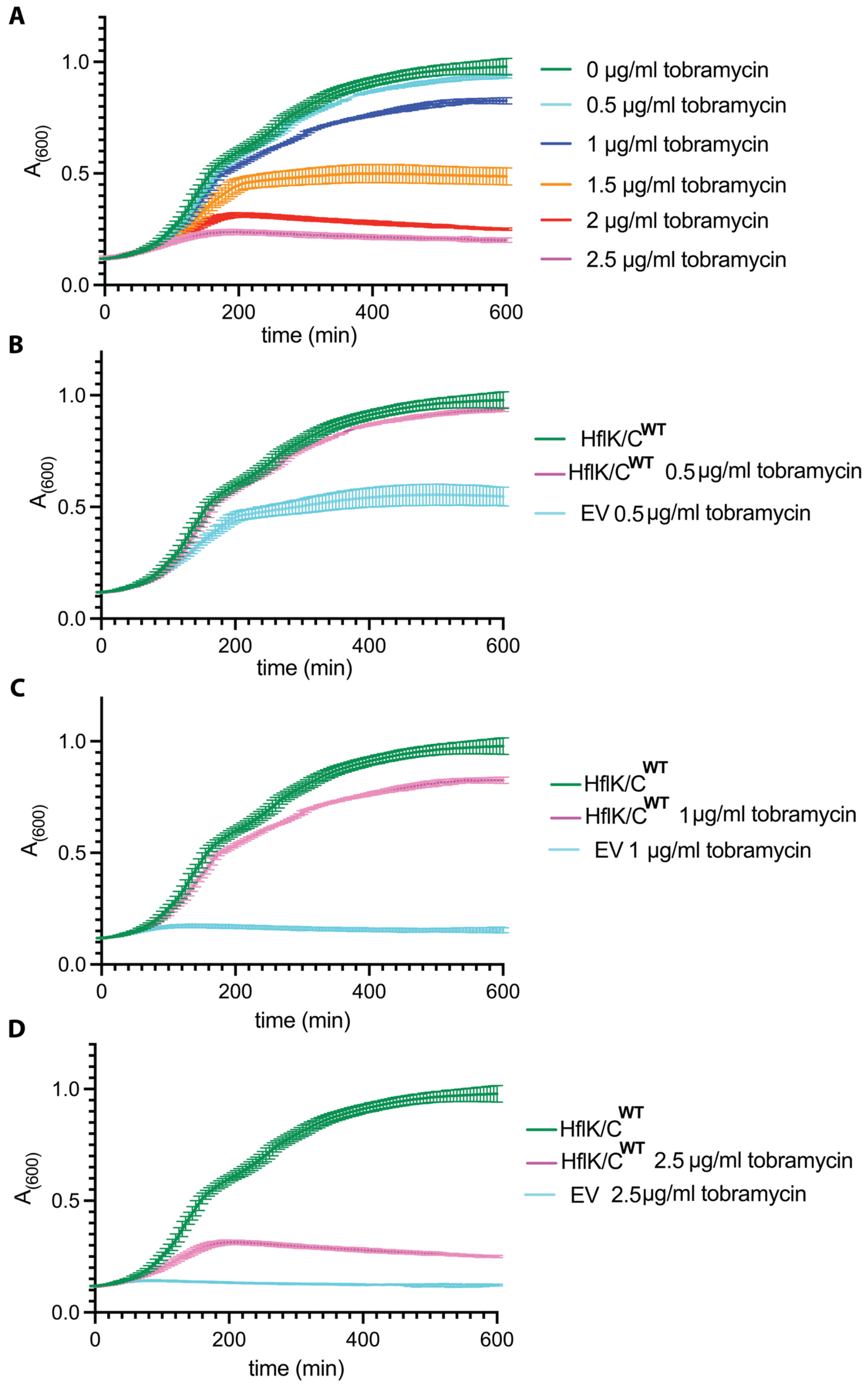


**Supplementary Figure 5. Inhibitory concentration of tobramycin for growth assays.** Growth assays were conducted using *E. coli* BL21 Δ*hflK/C* cells expressing wild-type HflK/C (HflK/C$^{WT}$) or an empty vector from a pPro24 plasmid under the control of a sodium propionate-inducible promoter at 37°C in LB medium. (**A**) Growth of cells expressing HflK/C$^{WT}$ across a range of tobramycin concentrations. (**B–D**) Growth rate comparisons between HflK/C$^{WT}$ and empty vector-expressing cells at 0-2.5 µg/mL tobramycin, respectively. Each growth assay was performed from a separate colony (n = 3 independent biological replicates), and data are presented as mean ± 1 SD.

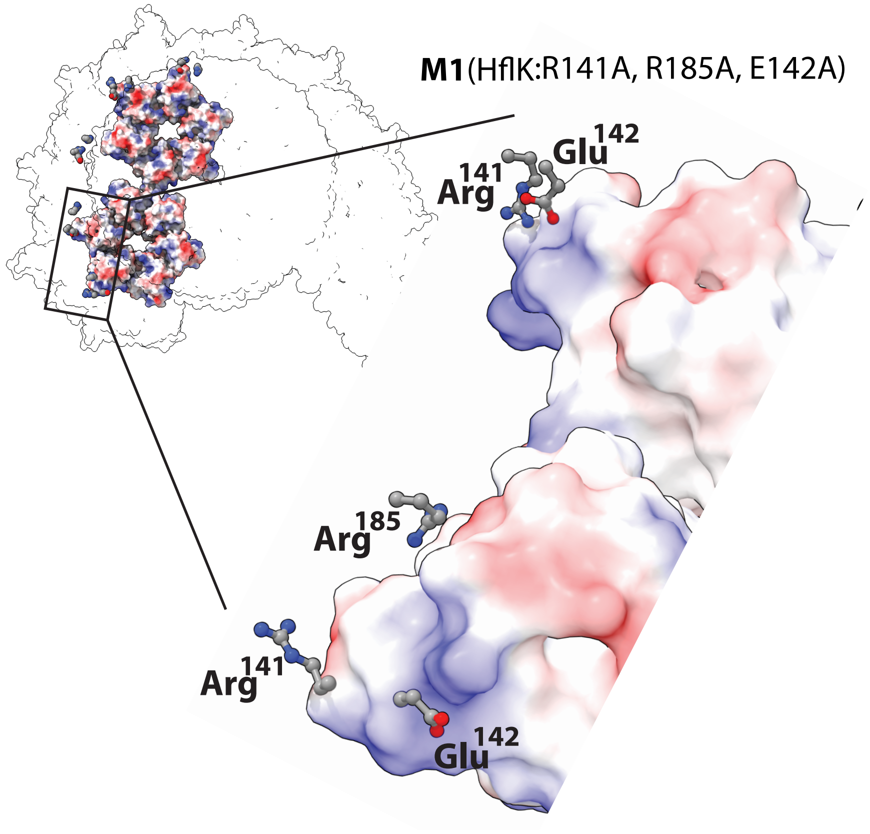


**Supplementary Figure 6. Mutational analysis of interface residues in the M1 HflK variant (R141A, E142A, R185A) within the HflK/C–FtsH complex.** Interface residues were identified based on the structural model of the FtsH•HflK/C assembly (PDB: 9CZ2).

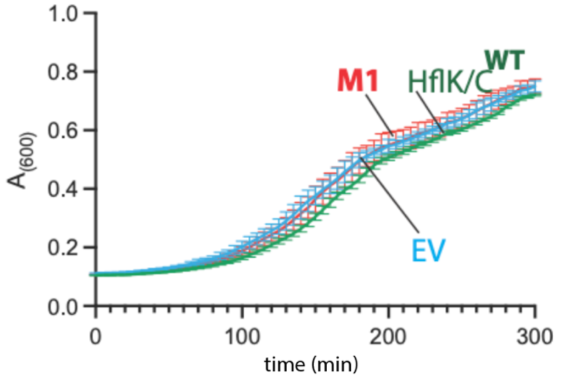


**Supplementary Figure 7. Growth assays for M1 mutants under a standard growth condition.** Growth assays were performed using *E. coli* BL21 Δ*hflK/C* cells expressing wild-type HflK/C(HflK/C$^{WT}$), **M1**, or an empty vector from a pPro24 plasmid under the control of a sodium propionate-inducible promoter at 37°C in LB medium. The growth assay was performed from a separate colony (n = 3 independent biological replicates), and data are presented as mean ± 1 SD.

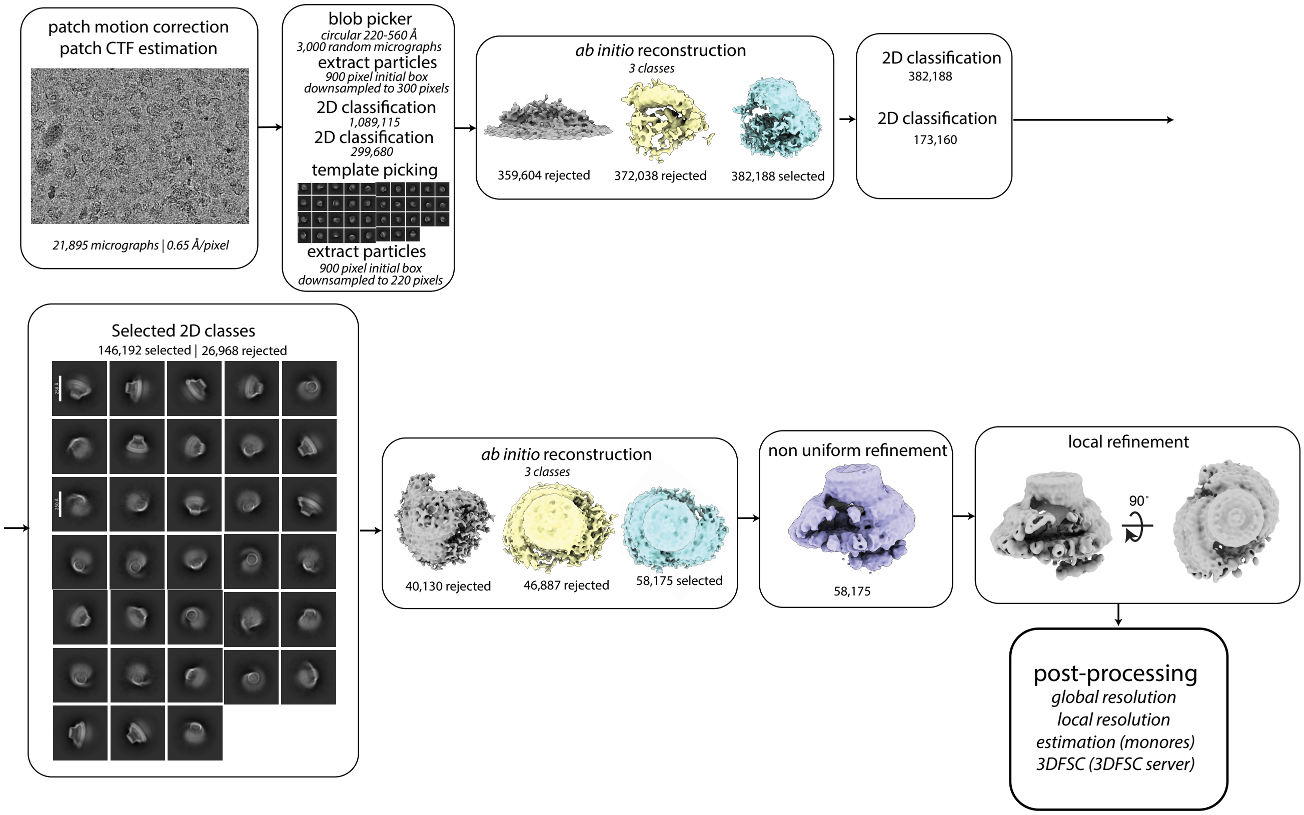


**Supplementary Figure 8. CryoSPARC processing workflow for the FtsH•HflK/C complex solubilized in DDM from cells treated with the aminoglycoside tobramycin.** The workflow outlines the cryo-EM data processing steps, including job names, job details, and any non-default parameters (italicized).

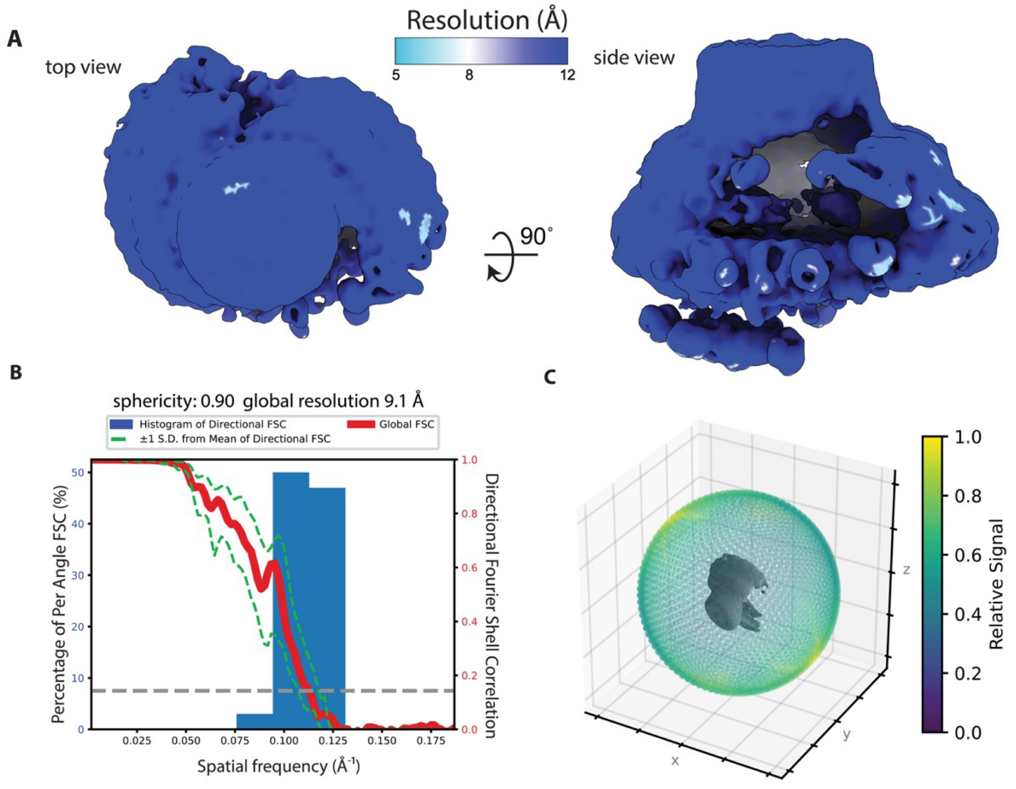


**Supplementary Figure 9. Estimates of resolution and angular sampling: DDM-solubilized FtsH•HflK/C complex extracted from the tobramycin treated cells**. **(A)** Maps colored by local resolution as estimated by the cryoSPARC implementation of monoRes. **(B)** Global resolution and directional resolution calculated by 3DFSC server (https://3dfsc.salk.edu). **(C)** Projection angle distribution estimated by cryoSPARC.

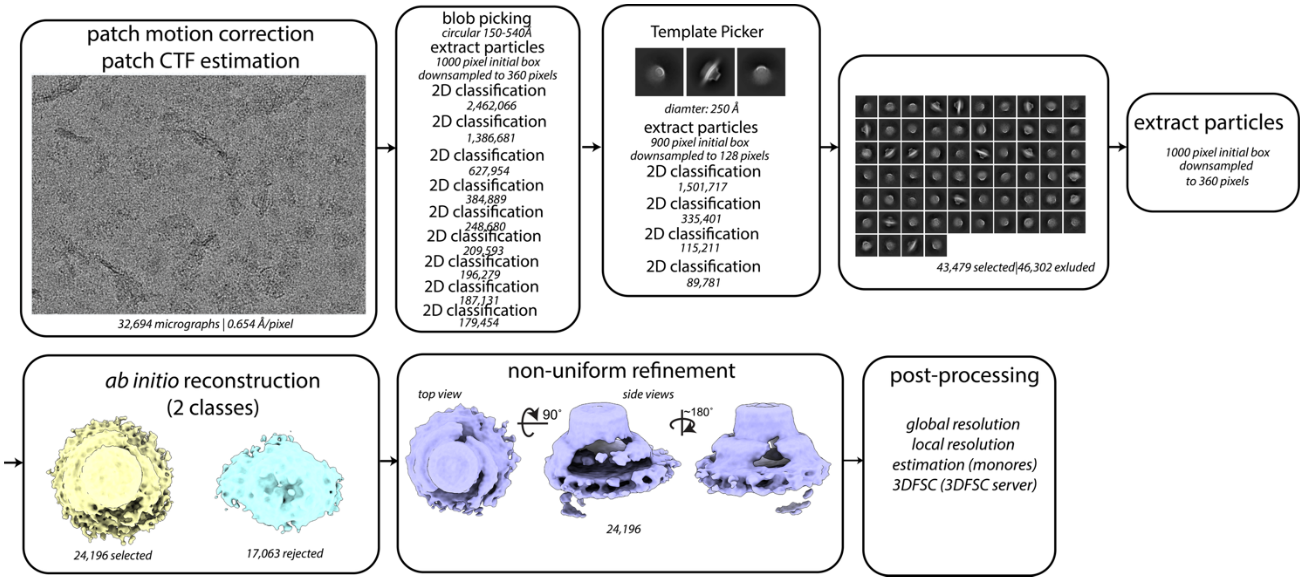


**Supplementary Figure 10. CryoSPARC processing workflow for the FtsH•HflK/C complex reconstituted in Carboxy-DIBMA from cells treated with the aminoglycoside tobramycin.** The workflow outlines the cryo-EM data processing steps, including job names, job details, and any non-default parameters (italicized).

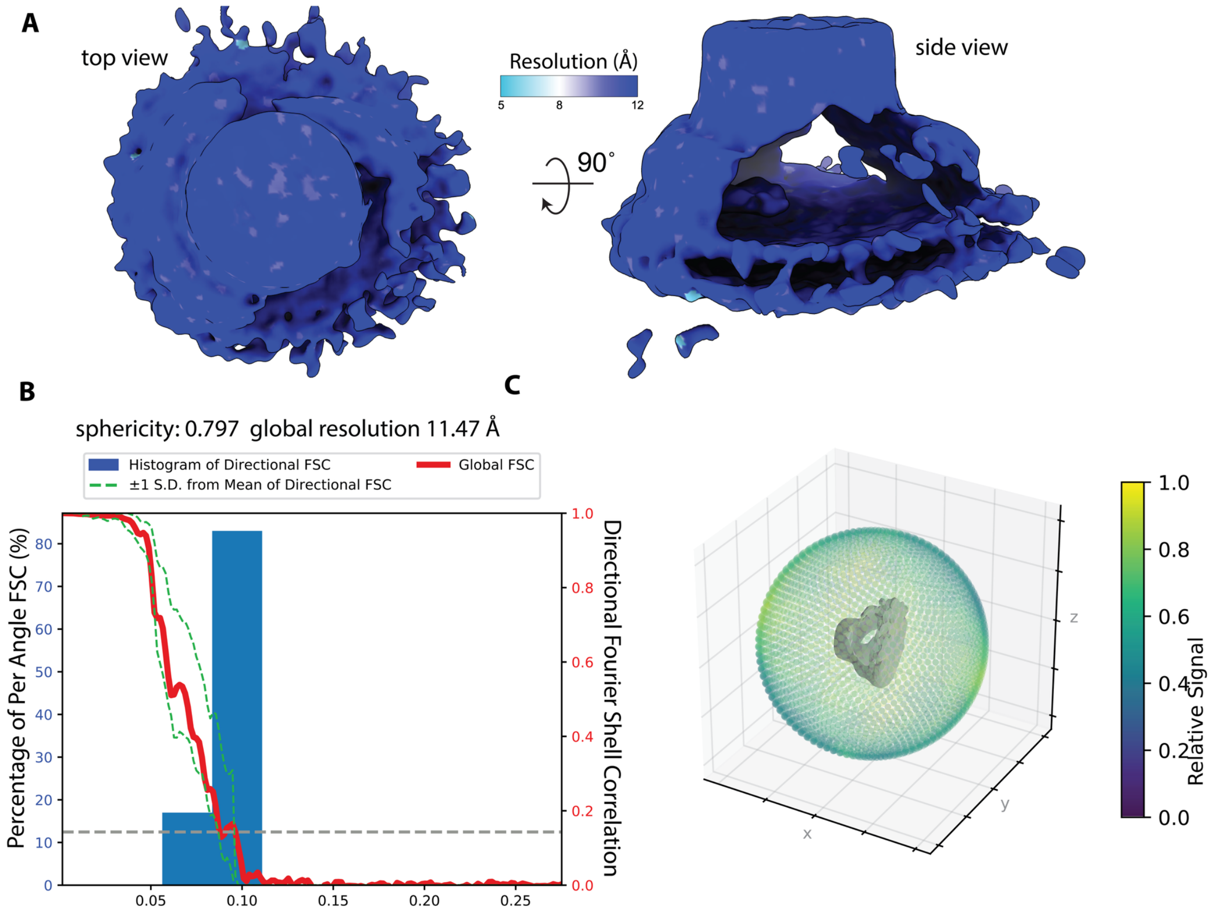


**Supplementary Figure 11. Estimates of resolution and angular sampling: Carboxy-DIBMA-extracted FtsH•HflK/C complex obtained from tobramycin-treated cells (map a**). **(A)** Maps colored by local resolution as estimated by the cryoSPARC implementation of monoRes. **(B)** Global resolution and directional resolution calculated by 3DFSC server (https://3dfsc.salk.edu). **(C)** Projection angle distribution estimated by cryoSPARC.
